# Dynamics of residual malaria transmission in Central Western Senegal: Mapping the breeding sites of *Anopheles gambiae* s. l

**DOI:** 10.1101/2020.07.13.200279

**Authors:** A. Ndiaye, E.A Niang, A. Niang-Diène, M. A. Nourdine, P C Sarr, L. Konaté, O. Faye, O. Gaye, O. Sy

**Author notes:** Corresponding authors: Ousmane Sy, Cheikh Anta Diop University of Dakar, Senegal E-mail: Ousmane Sy. Co-authors : Assane Ndiaye, El Hadji Amadou niang, Aminata Niang Diène, Mohamed Abderemane Nourdine, Pape Cheikh Sarr, Lassana Konaté, Ousmane Faye, Oumar Gaye.

## Abstract

Despite the use of several effective control interventions in the central western Senegal, residual malaria transmission still occurring in some hotspots. In order to better understand the factors associated with this situation to better tailor targeted control actions, it is critical to unravel environmental and geographical factors underlying the persistence of the disease in study hotspot villages. Hotspots villages were defined as those reporting more than six indigenous malaria cases during the previous year. A total of ten villages, including seven hotspots and three non-hotspots, were surveyed. All potential mosquito breeding sites identified in and around the tenth study villages were regularly monitored between 2013 and 2017. This monitoring concerned the presence of anophelines larvae and the collection of epidemiological, hydrogeological, topographical and biogeographical data. Throughout the study area, the number of larval breeding sites inventoried and monitored over the study period ranged from 50 to 62. They were higher, with no significant difference, in hotspot sites than in non-hotspot sites for each year of monitoring with 62.3% (56/62) in 2013, 90.9% (50/55) in 2014, 90.3% (56/62) in 2015 and 86% (43/50) in 2017 (Fisher’s exact test; p = 1). The Hotspot villages were mostly characterized by saline or moderately saline hydro-morphic and halomorphic soils allowing water retention and a suitable presence of potential larval breeding sites. Whereas non-hotspot villages are characterized mainly by a high proportion of extremely permeable sandy-textured soils due to their porosity, which reduces water retention. The annual number of confirmed malaria cases was corelated relatively with the frequency and extent of breeding sites. Malaria cases were much more higher in the hamlets located near to the breeding sites of *An. gambiae* s.l then gradually decreases with their remoteness. This study has shown that the dynamics of larval breeding sites by their longevity, stability, proximity to houses and their positiveness rate for the presence of *Anopheles* larvae could be a determining factor in the persistence of malaria hotspots in central western Senegal. The results of this study shed more light on the environmental factors underlying the residual transmission and should make it possible to better organize vector control interventions for malaria elimination in west-central Senegal.

## Introduction

Malaria is still a major parasitic public health problem worldwide, particularly in the sub-Saharan Africa. In 2019, the World Health Organization (WHO) estimates that 228 million cases of malaria occurred worldwide in 2018 with nearly 405,000 deaths, of which the main victims were pregnant women and children under 5 years. Noteworthy, up to 93% of the cases and 94% of the deaths still occurring in Africa, and the remaining cases and deaths are recorded mainly in the Southeast Asia and the Eastern Mediterranean regions [1].

In Senegal, malaria is endemic with seasonal recrudescence during the rainy season. In 2017, there were 395,706 confirmed malaria cases nation-wide, which have caused 284 deaths [2]. Nevertheless, despite this heavy burden, the disease incidence has declined since 2006 due to the massive increase of investments supporting both the prevention and the management of malaria cases. In the Central Western Senegal, specifically in the health districts of Mbour, Fatick and Bambey, in addition to the classical national strategies rolled out by the National Malaria Control Program (NMCP), Seasonal Malaria Chemo-prevention (SMC) was implemented between 2008 and 2011 followed by two targeted Indoor Residual Spraying (IRS) campaigns in 2013 and 2014 [3]. As a result of these different interventions, malaria is currently experiencing a very significant decline across the area, creating a new epidemiological profile with few villages where residual malaria transmission still occurring (hotspots villages) next to villages without any noticeable transmission [4]. This residual transmission may be due to environmental factors leading to the persistence of larval habitats. In some African countries, larval source management, a strategy which includes larviciding and breeding sites reduction, has been shown to be effective in reducing malaria transmission [5, 6, 7]. Information on the spatial distribution of mosquito breeding habitats is needed to enable monitoring and targeting of hotspot areas where vector populations are persistent. The development of Geographic Information Systems (GIS) mapping technology provides high-resolution maps of mosquito distribution or transmission risks, which could improve malaria surveillance system [8, 9].

This study was conducted to investigate the spatial variation and the underlying factors maintaining malaria transmission in some area versus others. More specifically, the main purpose of this study was to demonstrate the impact of anophelines larvae breeding sites location and functioning on the persistence of malaria transmission within hotspot villages in the central-western Senegal.

## Materials and methods

### Study area

The study was carried out in ten villages located in central western Senegal, spanning the three administrative departments of Mbour, Fatick and Bambey (Fig.1). The study area is located between the 13° and 15° 30 ’ Northern latitudes and the 13° and 16° 30’ Eastern longitudes, characterize by a flat relief, but sometime slightly uneven with altitudes rarely exceeding 20 m [10]. These low highlands are interspersed with valleys in which lowlands, ponds and creeks dried up very quickly during the dry season. As everywhere else in Senegal, the climate of the study area distinguishes a long dry season (7 to 8 months) and a very short rainy season (3 to 4 months). The major climatic features of the north Sudanese biogeographical region are the joint result of geographical and areological factors, with an annual rainfall averages ranging from 400 to 600 mm, unevenly distributed [11]. The monthly average temperatures are particularly high (above 30° C), between April to June. The soils type vary with terrain, bedrock, climate, and hydrography, with different agricultural aptitudes, evolving under the influence of human actions. In the study area, four types of soil with differences in texture, structure and organic matter content (sandy, clay, clay-sandy and salty soils) were identified. Located in the Sudan-Sahelian area, the vegetation is very diversified, and the geomorphological and pedological units combined with natural and human actions explain the density and diversity of the vegetal structure. Indeed, the vegetation is composed of treed, shrubby and herbaceous strata. The dominant economic activity in the area is agriculture, which mobilizes a large workforce, followed by cattle breeding, trade and crafts.

**Figure 1:**
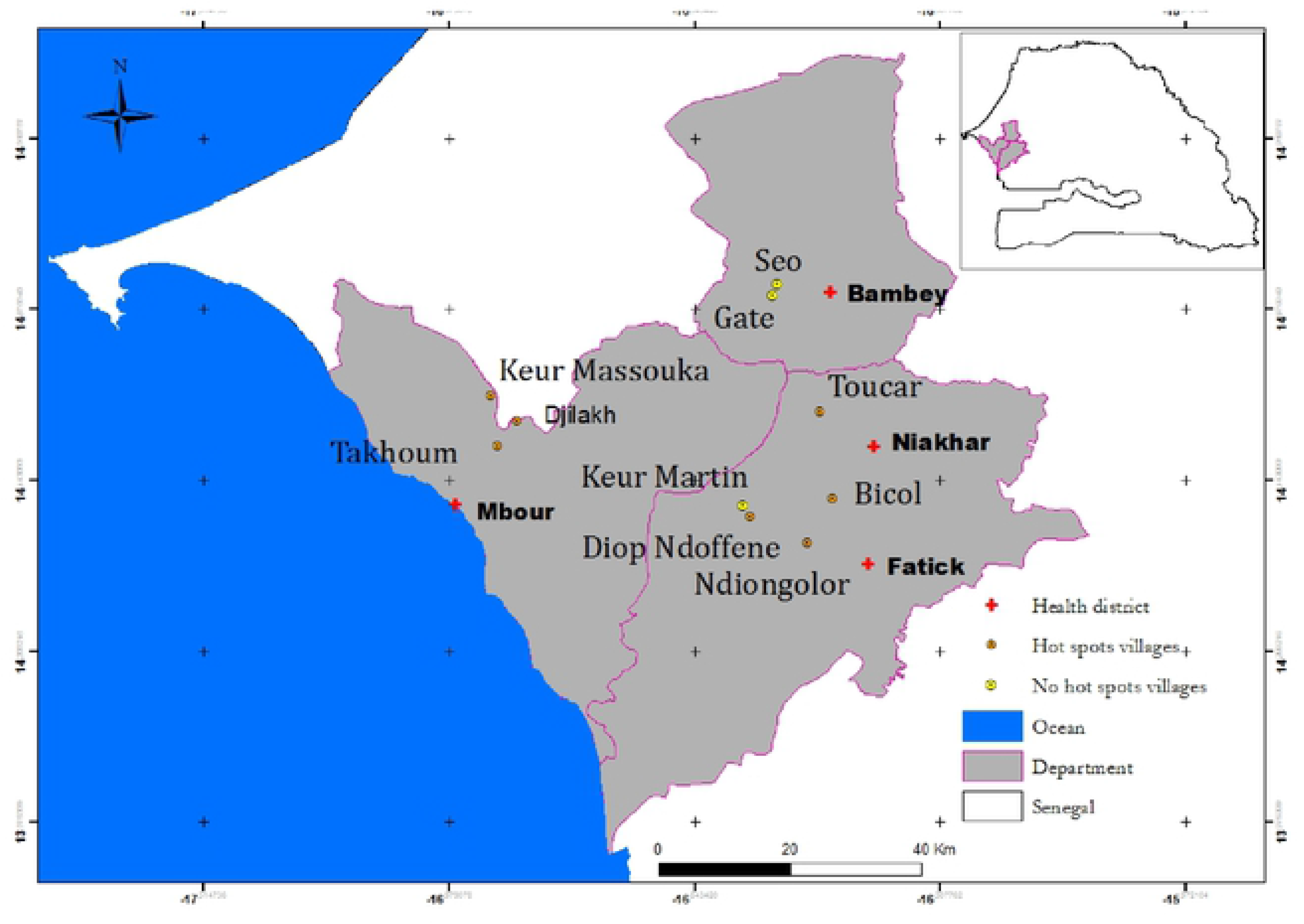
Study area (localization of health districts and epidemiological status of study villages)

### Selection of study villages

Overall ten villages were chosen based on their epidemiological status as hotspot (7 villages) or non-hotspot (3 villages). Hotspot villages were defined as those reporting more than 6 autochthonous malaria cases during the transmission period (from June to December) the previous year and reported by the health and demographic monitoring system implemented in the area since 2008 by the Department of Medical Parasitology of the Faculty of Medicine, Pharmacy and Stomatology of the University Cheikh Anta Diop of Dakar [4].

### Data collection

Data collected encompasses the geographical location and environmental factor prevailing in each of the selected study villages, as well as physical, chemical and biological parameters of encountered larval breeding.

### Characterization and mapping of larval breeding

The localization and the survey of the larval breeding sites within and around hotspot and non-hotspot villages were performed yearly during the raining season from 2013 to 2017. In each of the selected villages, all the encountered surface water bodies likely to harbor *Anopheles gambiae* s.l. larvae have been visually checked to assess the presence of larvae. Once the presence of larvae has been confirmed, useful information were recorded for each positive breeding site, including their coordinates using Geographic Positioning System (GPS) machine, Garmin International [12, 13]. The eggs deposition as well as larvae developmental rates were regularly monitored afterward. Therefore, all the encountered surface water bodies within and around the hotspot and non-hotspot villages which could potentially harbor anopheline larvae have been identified, mapped, then followed up over the time to assess their spatial distribution, their temporal functioning and their role in the maintenance of residual malaria transmission. Thematic maps were drawn using the thematic layers of the flora, the type of soil, the location of the water bodies (GPS coordinates), their distance to the nearest human dwellings, the type and putative origin of the water filling each water body and whether positive or not for Anopheles larvae using the “dipping” method.

### Mosquito larvae collection

Mosquito’s larvae were collected using the “dipping” method from each positive water body found within and close to each study village [14]. Upon collection, larval were sorted identified and separated as *Anopheles*, *Culex* or *Aedes* [15]. The remaining larvae and nymphs belonging to others mosquito’s genera were reared separately then identified using morphological identification keys [16]. The larval density within each positive breeding site has been calculated by counting the number of larvae per liter of water.

### Collection of morbidity data

Malaria morbidity was estimated from confirmed malaria cases using rapid diagnostic tests (RDTs) results for each of the tenth villages for a period lasting from the 1^st^ January 2013 to the 31^st^ December 2017. Data, including the age, the sex, RDT tests results (as positive or negative), the code and origin of patient, were retrieved from the consultation registers archived within the health structures of each study village. All sensitive information all patients have been made anonymous to comply the personal data protection requirement. Finally, morbidity data were collated with the geographical coordinates of all the districts and health structures visited for mapping purpose.

### Climatic data collection

One of the most important data when studying mosquito larval population is the rainfall to which the breeding sites are tributary. Rainfall data at the health district level were collected from the National Agency of Civil and Meteorological Aviation (ANACIM). Given the geographical location of the study villages, rainfall data were obtained from the rainfall stations located in the district of Mbour, Fatick and Bambey.

#### Data Processing and analysis

The geographic data of all GPS-tagged breeding sites and other information collected from the field were quality-checked before any analysis. Only data that were judged of good quality in terms of reliability of the source and data completeness were used for mapping and further analysis in Excel 2013, Epi info 7 and QGIS softwares. Data on larval breeding sites and human settlements were imported into the QGIS Desktop 2.2.0 mapping software to drawn maps and for spatial analysis.

## Results

### Relationship between the soil type and the frequency of breeding sites in hotspot and non-hotspot villages

The number of larval breeding sites inventoried and monitored between 2013 and 2017 ranged from 50 to 62 throughout the study area. They were higher, with no significant difference, in hotspot sites than in non-hotspot sites for each year of monitoring with 62.3% (56/62) in 2013, 90.9% (50/55) in 2014, 90.3% (56/62) in 2015 and 86% (43/50) in 2017 (Fisher’s exact test; p = 1). Moreover, across the study area, the occurrence and abundance of anopheline larval breeding sites were especially correlated to the rainfall and the nature of the soil. While in the non-host spot villages the soils were mainly halomorphic with more or less a clay texture, those in the hotspot areas were of tropical ferruginous types with little leaching with fine texture and high proportion of silt, and a fairly high content of clay, hence increasing their capacity for water retention. Moreover, in term of the level of the deposits at the immediate surroundings areas, hydromorphic soils with a higher rate of clay than the halomorphic ones were most often found. These types of soils were found mainly in the villages of Bicol, Ndiongolor and Toucar (Fig.1). They were also present in the north of the village of Keur Masssouka and throughout the eastern part of the villages of Djilakh and Takhoum Ndoundour. On the other hand, the halomorphic soils were characterized by the presence of a denatured surface and were more often connected to the ramifications of the inlets. These type of soils were found in the village of Diop Ndoffene and the non-hotspot village of Keur Martin. In the non-hotspot villages only few larval breeding, 11.7% (5/51,25) were found (Table 1), consisting mainly by borrow pits. Moreover, in the village of Gate, the anthropogenic breeding sites consisted of filled-up tire tracks left by heavy vehicles mired during the construction of the lateritic track connecting the village to the national road. These type of water bodies, characterize by a high proportion of sand-textured soils extremely permeable due to their porosity decreasing the water retention. They also have an unstable texture and dry up very quickly when the rains stop. These types of soils are very common in the villages of Gate and Seo, where no natural anopheline breeding site were found. Noteworthy, only two natural breeding sites were found across the surveyed villages in the non-hotspot area, precisely located in the village of Keur Martin (Fig.1).

**Table 1:** Number of breeding sites surveyed by village from 2013 to 2017

|  |  |  | Number of *An. gambiae* larval breeding sites per year |  |  |  |
| --- | --- | --- | --- | --- | --- | --- |
| Health districts | Villages | Epidemiological statut | 2013 | 2014 | 2015 | 2017 |
| Mbour | Djilakh | Hotspot | 9 | 9 | 9 | 5 |
|  | Takhoum Ndoundour |  | 2 | 2 | 2 | 2 |
|  | Keur Massouka |  | 5 | 5 | 5 | 3 |
| Fatick | Keur Martin | Non-hotspot | 2 | 2 | 2 | 4 |
|  | Diop Ndoffene | Hotspot | 7 | 7 | 7 | 7 |
|  | Bicol |  | 9 | 9 | 9 | 7 |
|  | Ndiongolor |  | 15 | 10 | 15 | 12 |
| Niakhar | Toucar | Hotspot | 9 | 8 | 9 | 7 |
| Bambey | Gate Diokoul 2 | Non-hotspot | 2 | 1 | 2 | 1 |
|  | Seo |  | 2 | 2 | 2 | 2 |
| Total in hotspot area |  |  | 56 | 50 | 56 | 43 |
| Total in non-hotspot area |  |  | 6 | 5 | 6 | 7 |

### Correlation between rainfall and vector breeding sites frequency

The rainfall significantly affects the production, availability, lasting and quality of anopheles larval breeding sites. In the study villages, the occurrence of breeding sites depended on the frequency and amount of rain. From 2013 to 2017, a total of 62 breeding sites were surveyed across all the study area (Table 1). The number and size of known and surveyed breeding sites varied between the beginning and the end of the rainy season. Overall there was a positive correlation (r = +0.089) between the rainfall and the proliferation of the breeding sites (Fig.2). In July, when the first rain were recorded in the area, only 67.7% (42/62) of the surveyed surface water body sites were filled up with water which size varying between 1.5 to 17,000 m^2^. In the middle of the rainy season, corresponding to the pick of rainfall between August and September, the number and size of the surface water bodies were the highest with 96% of the known breeding site filled up, their surfaces varying between 25 and more than 200000 m^2^. By the end of the rainy season in October, the proportion of the breeding sites dropped to 61% as well as their size varying between 3 and 25,000 m^2^. Unsurprisingly, the number and size of surface water bodies increased with the rainfall, but also according to the nature of the substrate.

**Figure 2:**
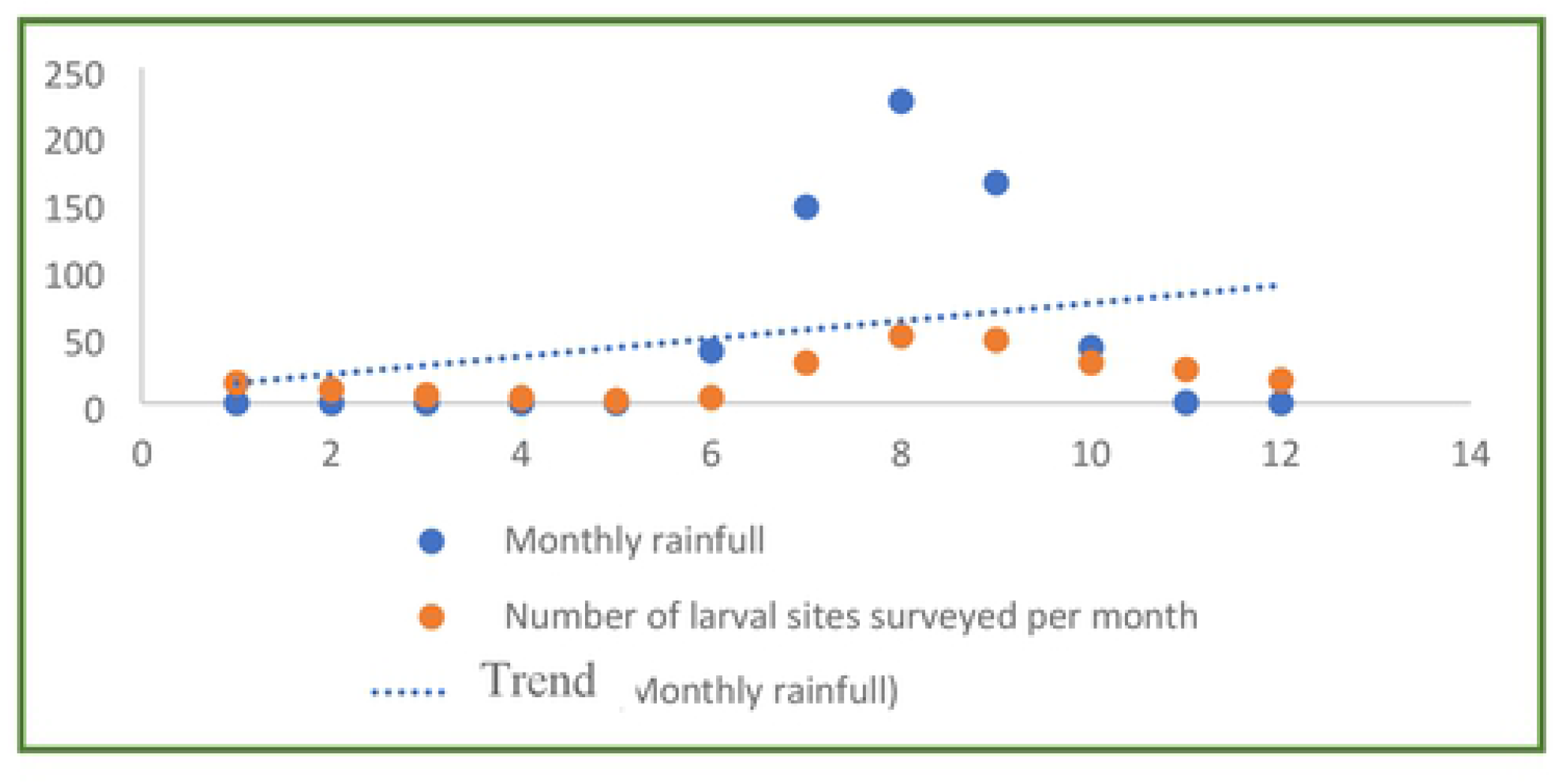
Evolution of the monthly average index of the frequency of *Anopheles gambiae* s.l. breeding sites according to monthly rainfall.

### Relationship between morbidity, frequency and size of the breeding sites of *An. gambiae* s.l. in the different epidemiological settings

Across the ten study villages, the number and size of *An. gambiae* larval breeding sites have varied over the study years. Overall the number of the breeding sites was 62 in 2013, 55 in 2014, 62 in 2015 and 50 in 2017 (Table 1). In the meantime, the overall malaria incidence varied with 0.109‰ in 2013, 0.064‰ in 2014, 0.142‰ in 2015 and 0.036‰ in 2017. In general, the annual number of malaria cases has followed relatively the frequency of larval breeding sites. It was higher in 2013 (26.1%) and in 2015 (35.2%) due to the significant rainfall recorded in the area for both years (Table1). The analysis of the frequency of confirmed malaria cases shows a similar evolution to that of the number of breeding sites surveyed. It was also noted that the highest number of malaria cases (93%) were recorded in the hotspot villages where the maximum number of breeding sites (50) were found. The cases were relatively low (7%) in the non-hotspot villages where the breeding sites were also found in lesser proportion (12) (Table 2). The drop of malaria cases in the hotspot villages of Bicol with (4.7%) and Diop Ndoffene with (2.1%) could be explained by the absence of morbidity data for the years 2013, 2014 and 2015 because of the loss of few consultations register (Table 2). In some villages, such as Keur Martin and Diop Ndoffene, the breeding sites correspond to large natural depressions of sizes up to over 200000 m^2^, most often temporary, filled by rainfall and overflowing seawater. These runoff inputs from marine or fluvial waters affect the neighbouring breeding sites level of salinity, which has repercussions on the presence, density and composition of anopheline larvae and therefore malaria morbidity.

**Table 2:** Number of confirmed malaria cases by village from 2013 to 2017

|  |  |  | Number of annual malaria cases |  |  |  |  |
| --- | --- | --- | --- | --- | --- | --- | --- |
| Health districts | Villages | Epidemiological statut | 2013 | 2014 | 2015 | 2016 | 2017 |
| Mbour | Djilakh | Hotspot | 35 | 14 | 53 | 19 | 4 |
|  | Takhoum Ndoundour |  | 12 | 12 | 21 | 4 | 5 |
|  | Keur Massouka |  | 20 | 5 | 11 | 3 | 1 |
| Fatick | Keur Martin | Non hotspot | 0 | 0 | 12 | 2 | 1 |
|  | Diop Ndoffene* | Hotspot | NA | NA | NA+5 | 3 | 1 |
|  | Bicol* |  | NA | NA | NA | 11 | 9 |
|  | Ndiongolor |  | 22 | 20 | 25 | 12 | 5 |
| Niakhar | Toucar | Hotspot | 14 | 12 | 16 | 5 | 9 |
| Bambey | Gate Diokoul 2 | Non hotspot | 5 | 1 | 4 | 2 | 1 |
|  | Seo |  | 1 | 0 | 0 | 0 | 0 |
| Total in hotspot area |  |  | 103 | 63 | 131 | 57 | 34 |
| Total in non-hotspot area |  |  | 6 | 1 | 16 | 4 | 2 |

\* Villages with missing data NA: Not Available

### Evolution of malaria cases according to larval breeding sites density in hotspots and non-hotspots villages

Larval breeding sites densities varied over the rainy season and impacted malaria evolution both in hotspots and non-hotspots villages (Fig.3). The strongest density was recorded in October and was mostly marked in hotspots villages with an average rate of 2 larvae per liter. It was twenty-five time lowest in non-hotspots villages, with an average rate of 0.08 larvae per liter. Overall malaria morbidity and vector breeding occurence were positively correlated (r^2^ = 0.63). Moreover, both parameters displayed similar trend over the time in the hotspot villages as well as in the non-hotspot villages. Overall at the beginning of the rainy season in July, the average *Anopheles gambiae* breeding sites found was 20.5%, rising to 53.2% in the middle of the rainy season in September, and 61.8% at the end of the rainy season in October (Fig.4). More than the half (51.87%) of the malaria cases recorded in the area in the zone occurred between the months of September and October, coinciding with the peak of the positivity of the breeding sites.

**Figure 3:**
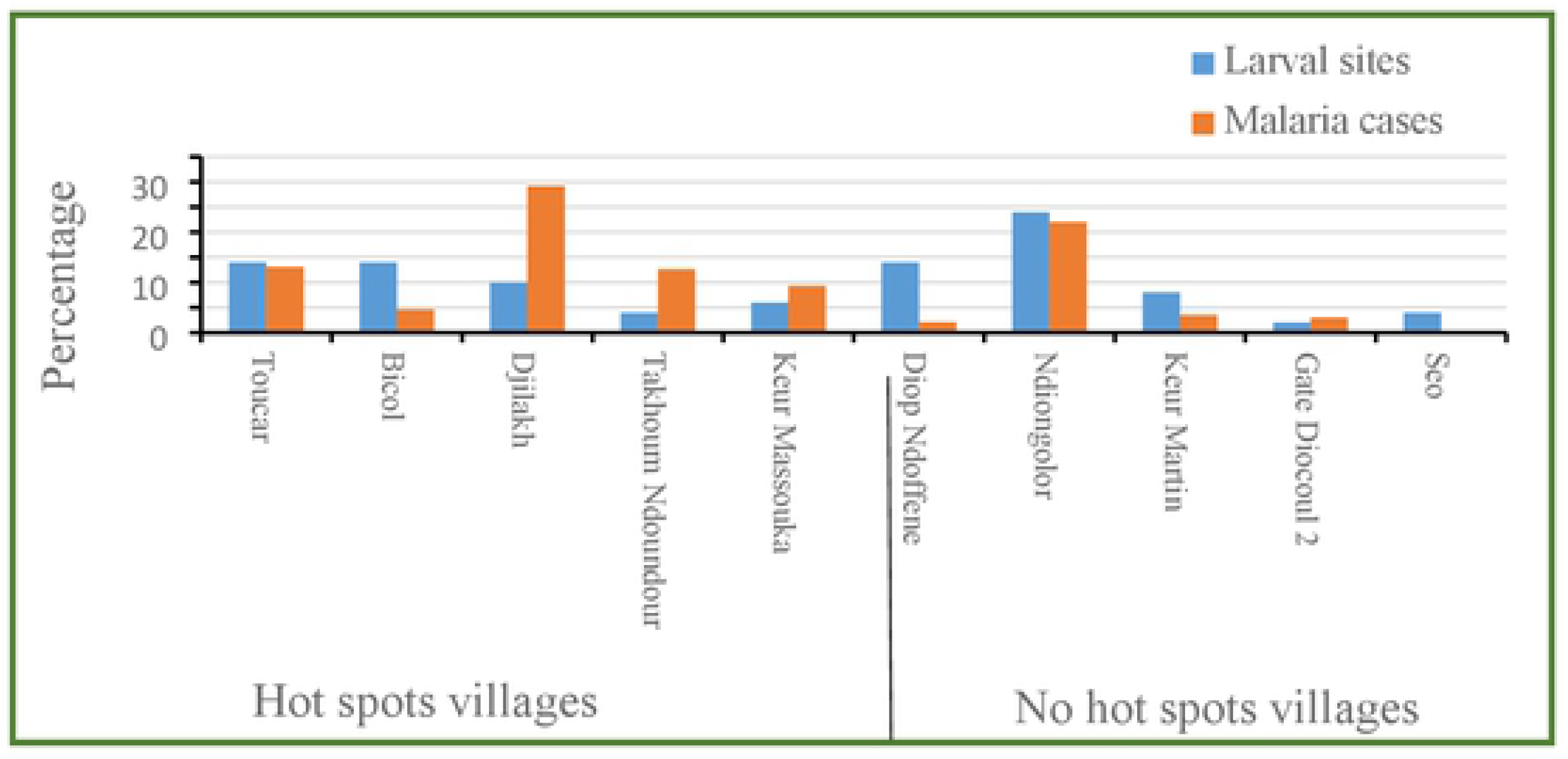
Relationship between morbidity and frequencies of *An. gambiae* sl breeding sites in hotspot and non-hotspot villages between 2013 and 2017

**Figure 4:**
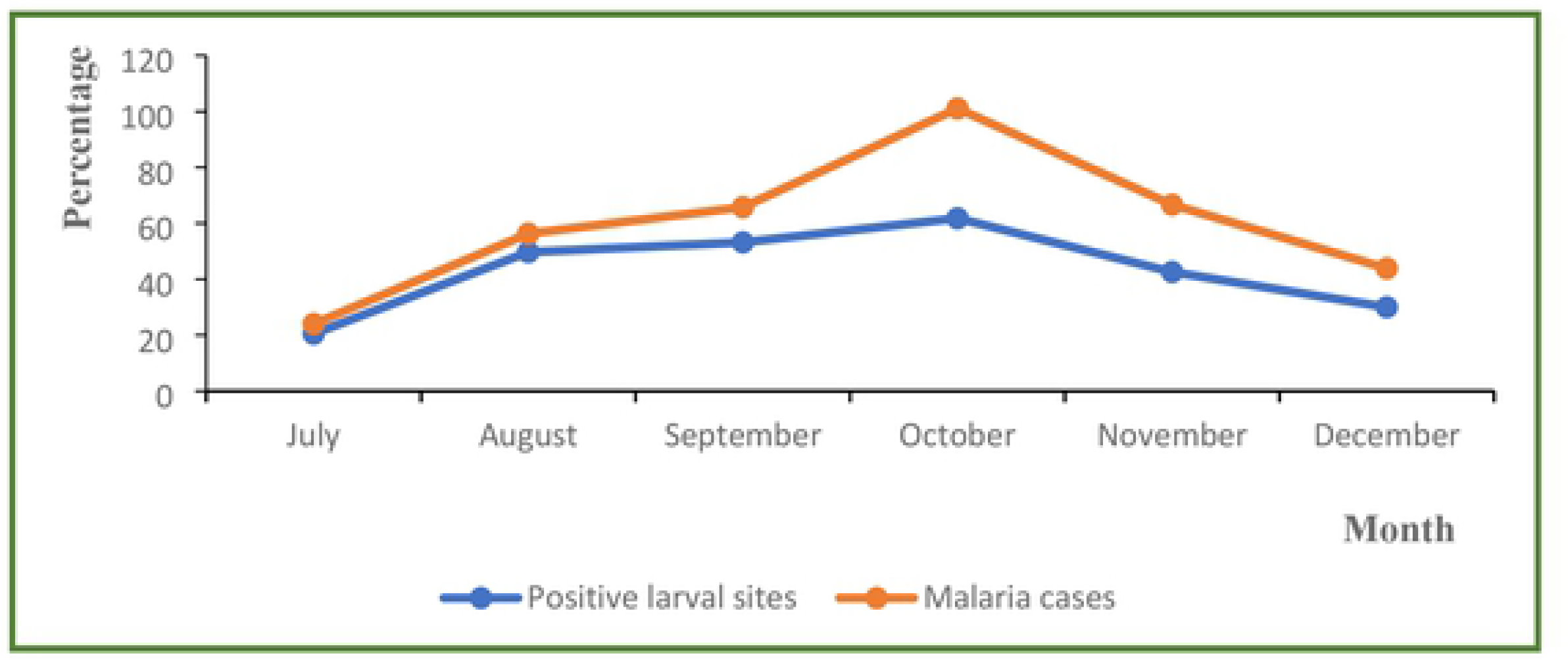
Relationship between monthly evolution of malaria cases and positivity of *An. gambiae* s.l. larval breeding sites between 2013 to 2017 in the ten study villages

### Variation of malaria morbidity according to the proximity of breeding sites and human dwellings: The case of the hotspot village of Djilakh

In the village of Djilakh in the district of Mbour, which is constituted by a group of hamlets, the analysis of morbidity data according to the location of *An. gambiae* s.l. larval breeding sites in relation to dwellings revealed three geographical areas displaying different level of the disease eco-epidemiology (Fig.5). Overall, malaria cases were significantly higher in the hamlets located the closest to breeding sites (< 500m) with 82.4% (103/125), followed by the hamlets located between 500 to 1000m, with 17.6% (22/125), and finally those farer away (≥1000m), where no cases were recorded during the study period (P < 0.05). The soil composition was different across the village, being mainly constituted by hydromorphic soils with a fairly high contain of clay, hence increasing its water retention capacity. Overall, malaria incidence was the relatively higher in the eastern part of the village where more breeding sites were found compared to the western part of the village. Looking at the soils’ composition, only the eastern part of the village consists of hydromorphic clay soils, while the rest of the study site consist of tropical ferruginous soils with little or no leaching (known locally as “dior” soils). These soils have a sandy texture and are very permeable hence their low water retention capacity. Under these conditions, the risk of malaria transmission remains relatively low.

**Figure 5:**
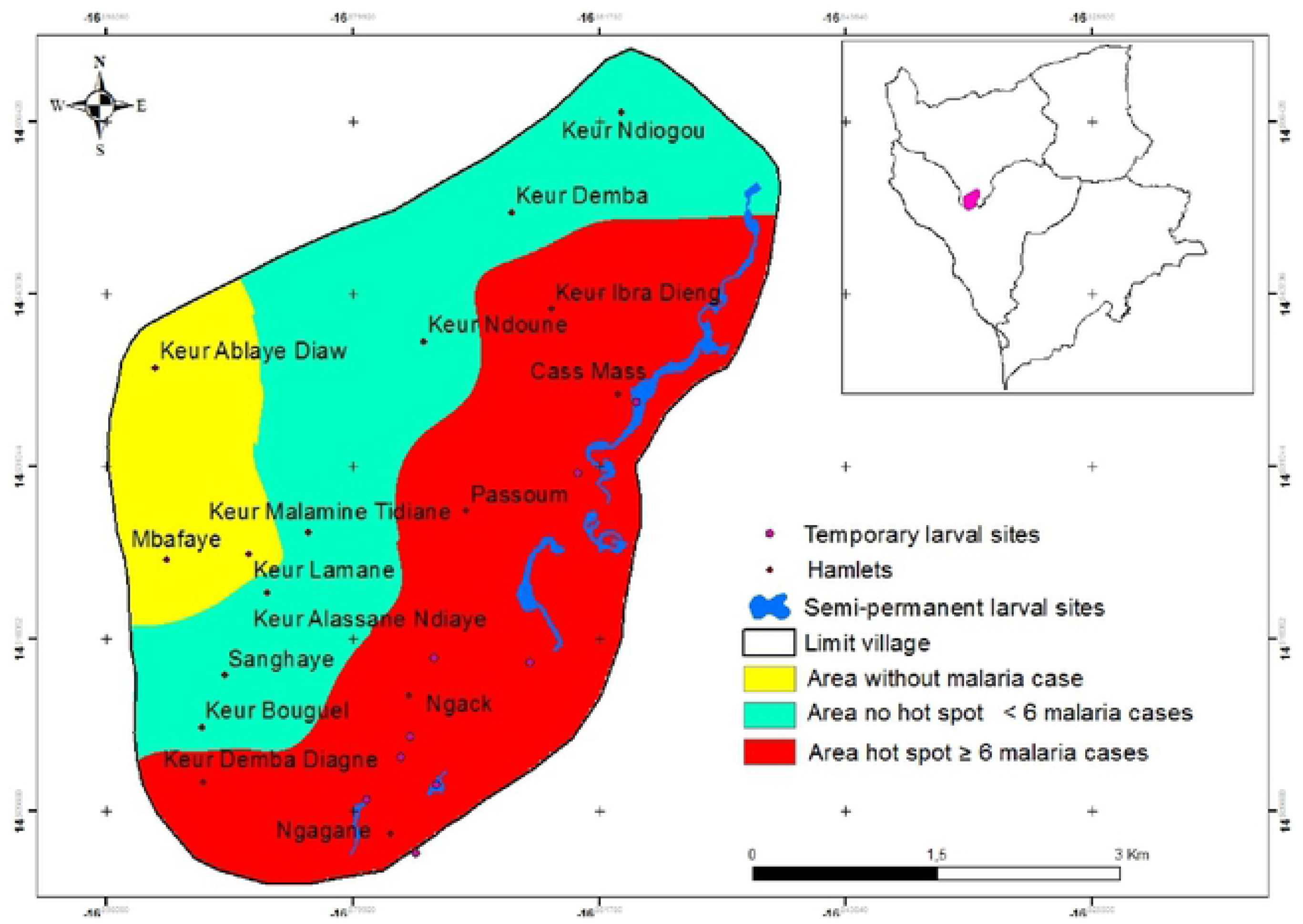
Malaria risk in Djilakh area: high risk of morbidity (in red), medium risk (in green) and risk-free (in yellow)

## Discussion

The conditions of malaria transmission are various, involving diverse ecological and environmental factors that may impact the disease epidemiology across the study area. The primordial biotope for the proliferation of mosquitoes results from the interaction between several environmental components such as water, the structure and composition of the soils and the vegetation associated with them [17, 18]. It is also important to note that humans play a key role in the influence of transmission and geographical distribution of the disease. Societies or human communities are not equal regarding the disease with sometime the creation of suitable or unfavorable conditions for the disease to spread [19].

The mineralogical composition of soils promotes the proliferation or not of putative mosquito breeding sites, especially when this combines with a significant humidity [20]. The creation of anophelines larval breeding sites is largely related to the nature of the soil substrates. This may explain why more breeding sites were found in the hotspot villages of Ndiongolor, Toucar, Bicol and Diop Ndoffene, where the soils consist mainly of hydromorphic and the saline halomorphic types, thus explaining the observed residual transmission. More specifically, hydromorphic and the halomorphic soils beside their capacity to retain flooding water over a long period of time, they are suitable for the larval development of some species of Anopheles such as *An. arabiensis* and *An. melas* depending on their salinity level. Indeed, in the localities where the hydromorphic soils were found and with surface water lasting longer, condition are favorable for the development of *An. gambiae* s.l. larvae, thus making them become places at high risk of malaria [21]. Conversely, in the non-hotspot villages of Gate Diocoul and Seo, the leached tropical ferruginous type of soils doesn’t allow surface water bodies to last for long period of time due to the significant infiltration of rainwater that the water table. Therefore fewer and very fast drying surface water points were found in those study sites, thus explaining lesser breeding sites recorded inside and around these villages as well as the lower levels of malaria morbidity.

The existence of suitable breeding sites lasting long enough for a complete larval development is fundamental to the life cycle of malaria vectors, and consequently to the transmission of the disease. The larval development is directly influenced by the volume of the water in breeding sites and their dynamics, the physical and chemical characteristics of the water, their solar exposure, the presence/absence and the type vegetation, occupation of the territory, but also by the average rainfall, water infiltration, evapotranspiration, and the water retention capacity of the soil etc…[22, 23, 24].

Most of anopheline breeding sites are filled during the raining season which thus corresponding to the malaria transmission season when vector populations pullulate. Across the study area, the transmission lasts for 5 to 6 months (from July to December) and is maintained by three main vectors species belonging to the Gambiae complex: *An. arabiensis, An. coluzzii* and *An. melas* [25, 26, 27]. As usually observed elsewhere across the country, malaria incidence increased progressively right after the first rains with a peak at the middle of the rainy season toward the month of October, while the peak of the rain was recorded earlier between August and September. This led to the proliferation of surface water collections suitable for the anopheline larval development [28]. The lag time between the peak of rainfall and the peak of transmission as observed here. This is a classical situation due to the fact that when rains increase and are recurrent, as observed in the months of August and September, breeding site become constantly flooded and the leaching of the breeding sites cause the death of immature larval stages [29]. Then when the dynamics of rainfall become more spaced, the breeding sites become more stable from September toward October and more productive thus changing the disease transmission dynamic. Indeed, this study has shown that the peak of morbidity follows with two months delay the peak of rains. The study also confirms previous observations in the area reporting a lag time of two months between the maximum rainfall and the peaks of malaria-related deaths [30]. Indeed, the larval development and abundance of malaria vector are closely related to the presence and productivity of breeding sites, which for the members of the *An. gambiae* complex are tributary of the dynamics of rainfall [31]. Having clearly demonstrated the link between malaria morbidity and the number of productive breeding sites in the study villages, the rainfall deficit observed in October 2014 explain the scarcity of breeding sites and consequently the lower malaria vectors density the same across the study area. Indeed, the large majority of known breeding sites dried up very quickly due the decrease of rains toward the end of September 2014 [25]. This resulted consequently with the drop of malaria morbidity as observed the same year, with a reduction of 45 confirmed malaria cases compared to the previous year (2013).

Among factors highly influencing the abundance of anopheline mosquitoes, beside the thermal and hygrometric, physical and chemical parameters of the breeding sites water are of the highest importance. Each mosquito species has its specific requirement for water properties for an optimal development during the larval stages. However, the physical and chemical parameters of the water vary over the time and with the rainfall dynamic but become more stable toward the end of the rainy season with the breeding site less diluted by recurrent rainfalls, thus favoring the proliferation of mosquitoes [32]. Indeed, in the Sudanese zone of West Africa, the peak of transmission is usually recorded towards the end of the rainy season when the humidity remains high enough while the rains are less frequent, no longer disturbing the breeding sites parameters, including their physical and chemical factors [33, 34].

The analysis of the breeding site location to human dwellings showed that malaria cases decreases with distance [35, 36]. For instance, in the village of Djilakh, the number of new malaria cases was much more higher in the hamlets nearby the breeding sites and where the soil type is more favorable for rainwater retention. The number of cases decreased gradually with distance. Thus, the areas most exposed to the risk of malaria transmission can be considered as those close to the main water retention areas and the most remote neighborhoods with soils that retain less water are the least exposed [21, 37]

## Conclusion

The results of this study showed that the dynamics of anopheles mosquitoes breeding sites with regard to their abundance, stability and proximity to houses strongly influence the epidemiology and the persistence of malaria transmission in hotspots villages in central western Senegal. A good management strategy of larval breeding sites in hotspot villages, aiming at reducing the number of mosquito immature stages could be integrated as an additional measure in this area where almost all functioning breeding sites are known, to further reduce transmission and achieve malaria elimination goal in central western Senegal.

## Authors contribution

Conceptualization: Ousmane Faye, Aminata Niang Diène, Lassana Konaté, Oumar Gaye, Ousmane Sy, Assane Ndiaye.

Investigation: Ousmane Sy, Assane Ndiaye, Pape Cheikh Sarr, Mouhamed Abderehman Nourdine.

Supervision: Ousmane Faye, Oumar Gaye, Aminata Niang Diène, Lassana Konaté, Ousmane Sy, Elhadji Amadou Niang, Assane Ndiaye.

Validation: Ousmane Faye, Aminata Niang Diène, Lassana Konaté and Oumar Gaye

Writing – review & editing: Ousmane Faye, Oumar Gaye, Lassana konaté, Ousmane Sy, Elhadji Amadou Niang, Aminata Niang Diène, Assane Ndiaye, Pape Cheikh Sarr, Mouhamed Abdereman Nourdine.

## Ethics approval and consent to participate

This study was approved by the National Ethics Committee of Senegal.

## Ethics approval and consent to participate

This study was approved by the Ethics Committee of University Cheikh Anta Diop of Dakar, Senegal.

